# A transmissible RNA pathway in honey bees

**DOI:** 10.1101/299800

**Authors:** Eyal Maori, Yael Garbian, Vered Kunik, Rita Mozes-Koch, Osnat Malka, Haim Kalev, Niv Sabath, Ilan Sela, Sharoni Shafir

**Affiliations:** Department of Veterinary Medicine, University of Cambridge, Madingley Road, Cambridge, CB3 0ES, United Kingdom; The Hebrew University of Jerusalem, The Robert H. Smith Faculty of Agriculture, Food and Environment, Rehovot, Israel; Vered Kunik – bioinformatics consulting, 12 Hailanot street, Gat-Rimon, 4992000, Israel; Department of Biochemistry, Rappaport Faculty of Medicine, Technion-Israel Institute of Technology, Haifa 31096, Israel

## Abstract

One of the characteristics of RNA interference (RNAi) is systemic spread of the silencing signal among cells and tissues throughout the organism. Systemic RNAi, initiated by double-stranded RNA (dsRNA) ingestion, has been reported in diverse invertebrates, including honey bees, demonstrating environmental RNA uptake that undermines homologous gene expression. However, the question why any organism would take up RNA from the environment has remained largely unanswered. Here, we report on horizontal RNA flow among honey bees mediated by secretion and ingestion of worker and royal jelly diets. We show that ingested dsRNA spreads through the bee’s hemolymph associated with a protein complex. The systemic dsRNA is secreted with the jelly and delivered to larvae via ingestion. Furthermore, we demonstrate that transmission of jelly-secreted dsRNA to larvae is biologically active and triggers gene knockdown that lasts into adulthood. Finally, RNA extracted from worker and royal jellies harbor differential naturally occurring RNA populations. Some of these RNAs corresponded to honey bee protein coding genes, transposable elements, non-coding RNA as well as bacteria, fungi and viruses. These results reveal an inherent property of honey bees to share RNA among individuals and generations. Thus, our findings suggest a transmissible RNA pathway, playing a role in social immunity and epigenetic signaling between honey bees and potentially among other closely interacting organisms.

**SIGNIFICANCE:** Honey bees are eusocial insects, living in a colony that is often described as a superorganism. RNA mobility among cells of an organism has been documented in plants and animals. Here we show that RNA spreads further in honey bees, and is horizontally transferred between individuals and across generations. We found that honey bees share biologically active RNA through secretion and ingestion of worker and royal jellies. Such RNA initiates RNA interference, which is a known defense mechanism against viral infection. Furthermore, we characterized diverse RNA profiles of worker and royal jelly, including fragmented viral RNA. Our findings demonstrate a transmissible RNA pathway with potential roles in social immunity and epigenetic signaling among members of the hive.

## Introduction

In eukaryotes, sequence-specific gene silencing pathways, generally termed RNA interference (RNAi), are induced and maintained by the presence of double stranded RNA (dsRNA) [1]. Through processing of based-paired RNA into small RNAs, these mechanisms regulate gene expression in both co-transcriptional and post-transcriptional levels [2]. While RNA-mediated nascent transcript destabilization and heterochromatin remodeling inhibits gene transcription, post-transcriptional gene silencing down regulates gene expression through guiding target RNA degradation or repression of translation [3,4].

RNAi could be divided into cell-autonomous and non-cell autonomous [5]. In cell-autonomous RNAi, silencing is restricted to cells that produce or were exposed to the dsRNA trigger. Initiation of local RNAi could develop in some organisms into a non-cell autonomous silencing signal, affecting cells and tissues which originally did not generate or were not introduced to dsRNA [6,7]. The mechanisms that facilitate RNA export from donor cells, extracellular spread and import into acceptor cells are not fully elucidated, but under ongoing investigation in diverse biological systems.

In 1998, Timmons and Fire were the first to report on gene silencing triggered by environmentally acquired dsRNA [8]. In essence, their finding represents a form of horizontal regulatory RNA transfer. To date, susceptibility to environmental RNAi has been established in fungus and animals from different phyla including Nematodes, Plathelminthes and Arthropods [reviewed in [9]]. Environmental RNAi experiments mostly utilize bacterially expressed- or in vitro synthesized dsRNA through ingestion, suggesting that dietary consumption is an effective RNA uptake pathway. Further supporting this, potent RNAi transmission from transgenic dsRNA-expressing plants to invertebrate herbivores has been widely reported [10,11]. Accordingly, host to-parasite RNAi transfer (commonly termed “host-induced gene silencing”; HIGS) has been applied to agriculture in recent years, demonstrating a potential practical strategy to control various pest and viral related diseases [12,13].

One of the main questions in the field of environmental RNAi is whether natural and functional RNA transfer among organisms occurs. Recently, transmission of parasitic nematode-derived miRNA to its mammalian host has been shown to compromise immunity of infected mice [14]. Similarly, pathogenic fungi exploit mobile small RNA signals to modulate via RNAi plant immune responses [15]. Reciprocally, arabidopsis plants secrete and transfer vesicles-containing small RNAs that could suppress virulence fungal genes [16]. Furthermore, ingestion of pollen-derived plant miRNA induces worker bee sterility in a sequence specific manner [17]. Interestingly, while the aforementioned examples provide evidence that some organisms acquire, and are affected by foreign regulatory RNA, it is still puzzling why would they allow it? This evolutionary maintained susceptibility to non-self regulatory RNA is intriguing in light of the fact that the most well-known transmissible RNA are viruses.

The honey bee (*Apis mellifera*) has been established as a model to study various disciplines in biology. One of the remarkable characteristics of honey bees is their environmentally-mediated phenotypic plasticity [18]. Female bee larvae can either develop into worker or queen; two castes with distinct morphology, physiology, reproductive capability, life span and behavior. This developmental flexibility of genetically identical individuals is driven by differential diet consumption. A larva fed exclusively on royal jelly will develop into a queen, whereas larva fed on worker jelly will develop into a worker [19]. In other words, nutritional differences trigger one genome to direct two distinct phenotypic outputs in honey bees.

The honey bee caste determination is driven by the interplay of food quantity and quality, environmental conditions and gene expression [20,21]. Epigenetic regulation has been suggested to play a role in the honey bee’s clonal phenotypic variation. Knockdown of DNA methyl-transferase 3 (dnmt3), a key enzyme in DNA methylation process, has a royal jelly-like effect, and semi-queen workers emerge with fully developed ovaries and a transcriptome profile similar to naturally reared queens [22]. Moreover, the methylation imprint varies between the brain’s DNA of workers and queens, demonstrating unique epigenetic profiles among workers and queens [23]. Consistently, (E)-10-hydroxy-2-decenoic acid (10HDA), a fatty acid that comprises up to 5% of the royal jelly, has been characterized as a histone deacetylase inhibitor (HDACi) [24]. Nonetheless, although the general involvement of epigenetics has been established, it is still not clear how caste-specific DNA methylation marking is directed.

Previously, we reported on a RNAi-based ingestion system for the control of Israeli Acute Paralysis Virus (IAPV) disease in honey bees [25]. Field trials, in which this environmental RNAi system was employed, indicated that the colony performance of virus-inoculated hives deteriorated following virus infection in control hives, whereas that of dsRNA-treated hives remained strong [26]. Interestingly, dsRNA-treated hives also produced more honey, when the main honey flow was 3-4 months after the last dsRNA treatment. By that time, most of the originally treated bees would have been replaced by new generations. Honey bee viruses can be transmitted among individuals in the hive both horizontally and vertically [27]. It was therefore expected that hives would become virus-affected once the new generation gradually replaced the previous, dsRNA-treated one. Potential persistence of disease protection raised the question whether treated honey bees may serve as vectors for RNA.

Following this hypothesis, here we show that environmentally consumed dsRNA in honey bees is up-taken from the digestive system and systemically spread through the hemolymph associated with a protein complex. Then, these RNAi carrier bees transfer silencing-triggering molecules to the next generation via dsRNA secretion into the jelly. Moreover, we demonstrate that jelly-secreted dsRNA is biologically active and triggers a long lasting silencing effect in the recipient generation. Finally, we characterize diverse naturally occurring endogenous and exogenous RNA populations in royal and worker jellies. These findings demonstrate an environmentally mediated transmissible RNA in honey bees.

## Results

### Intake of ingested dsRNA into the honey bee hemolymph

Adult honey bees exchange food via trophallaxis and all bees in a hive are regarded as having a “shared stomach” [18]. Consumed but pre-digested dsRNA may thus be distributed directly among adult bees. However, nurse honey bees nourish the young larvae for the first three days with a processed diet secreted from the mandibular and hypopharyngeal food glands, the worker and royal jellies [19]. Queens are exclusively fed on royal jelly their entire life. In order to transmit RNAi-triggering molecules to the next generation, the ingested dsRNA has to spread systemically and reach the food glands, and then be secreted in the jellies. Therefore, we first attempted to test whether ingested dsRNA occurs in the honey bee’s circulatory system, where it can systemically spread. To that end, we applied Digoxigenin (DIG)-labeled dsRNA (dsRNA*) for the direct detection of ingested dsRNA.

Adult worker bees were immobilized and fed on dsRNA* or mock solutions directly to their glossa in order to avoid cuticle contamination. Five hours post feeding, we extracted hemolymph of bees and tested for the presence of dsRNA* in whole raw hemolymph extracts. A labeled band corresponding to the purified dsRNA* size (430bp) was detected in the hemolymph of treated bees (Figure 1A), demonstrating uptake of full length dsRNA from the digestive system to the circulatory system.

**Figure 1.**
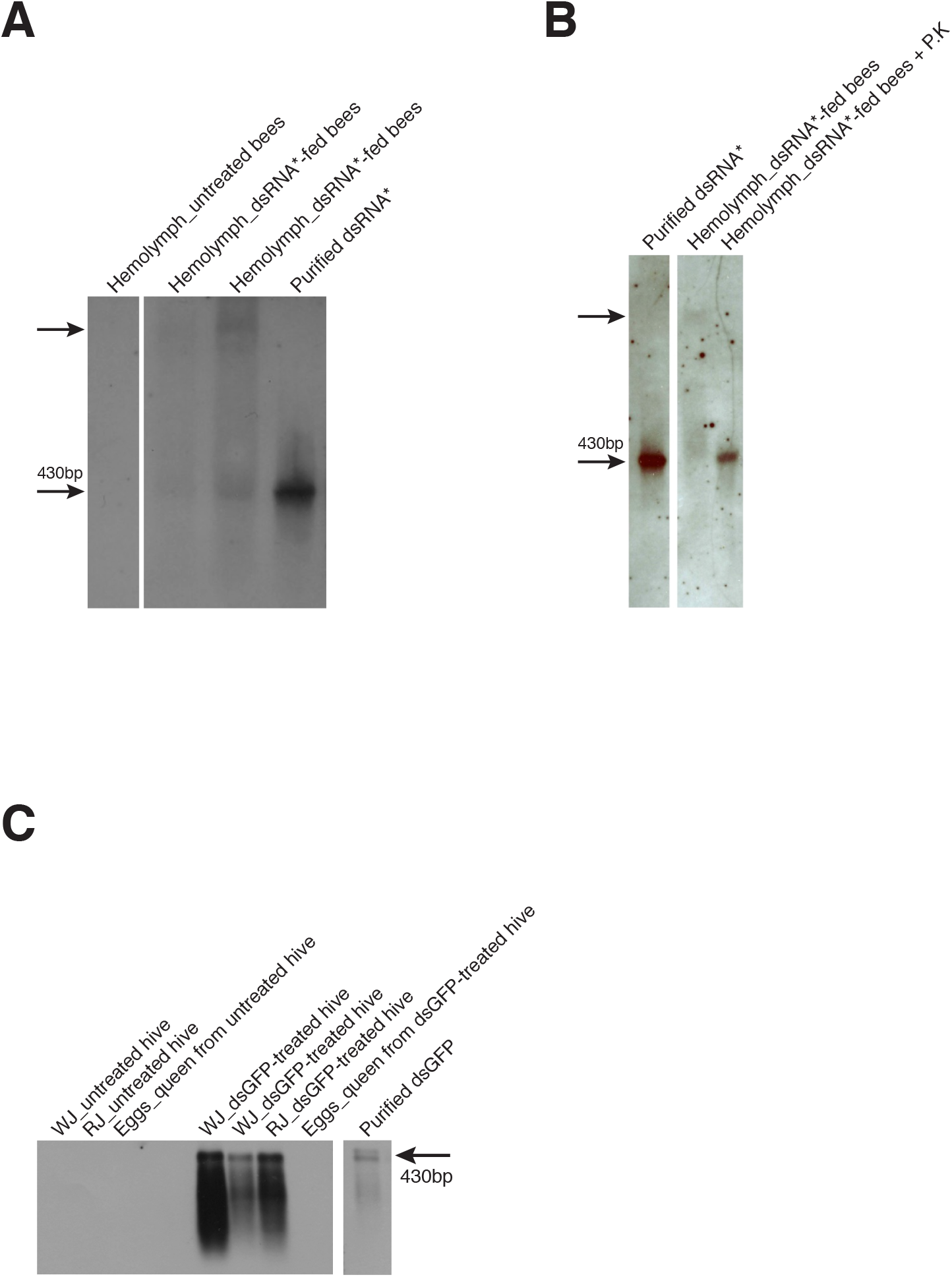
**A)** Presence of ingested dsRNA in the hemolymph. Probe-free Northern blot analysis performed on pooled raw hemolymph extracts (ten bees per pool, 10 μl per well). Raw hemolymph extracts were collected from bees that were fed on 50% sucrose solution (w/w) containing DIG-labeled dsRNA-GFP, or sucrose solution only. The 430 bp band represents free full-length dsRNA. **B)** Association of ingested-dsRNA with a protein complex in the hemolymph. Probe-free Northern blot analysis performed on untreated or protease K-treated pooled raw hemolymph extracts (ten bees per pool, 10 μl per well). The hemolymph in both treatments was derived from the same hemolymph sample. **C)** Occurrence of ingested dsRNA in the larval diets and newly laid eggs from dsRNA-GFP treated and untreated minihives. Northern blot analysis of 1 μg total RNA extracted from worker jelly (**WJ)**, royal jelly (RJ) and eggs. Eggs were laid in untreated minihives by queens that were transferred from dsRNA-treated or untreated minihives. Purified dsRNA-GFP was used as a positive control and a size marker for full-length dsRNA.

Interestingly, two additional dsRNA* bands of higher molecular weight (corresponding approximately to 2.5 and 4 Kbp) were also observed in raw hemolymph extracts (Figure 1A). Two scenarios can be drawn to explain the appearance of high molecular weight labeled RNAs: (i) recombination between dsRNA* and other RNAs; or (ii) association of the dsRNA* with other components of the hemolymph. Following Proteinase K digestion of the hemolymph extract the size of the higher bands shifted back to the expected 430 bp, indicating that the higher compounds were complexes of ca. full length dsRNA and hemolymph protein(s) (Figure 1B). It is worth noting that no processed dsRNA forms could be detected in the hemolymph extracts tested.

### Presence of ingested dsRNA in the worker and royal jelly

The honey bee’s hemolymph is a complex fluid mixture of immune cells (haemocytes), lipids, carbohydrates, nucleic acids, hormones and proteins [28]. The finding of ingested dsRNA-protein complexes in the hemolymph indicates a possible active mechanism for uptake and translocation of dsRNA through the honey bee’s circulatory system and possible spread to the jelly producing glands. If this happens, mobile dsRNA may be present in the jelly. Therefore, we next tested whether ingested dsRNA is secreted in the diet of worker- and queen-destined larvae.

Minihives with ca. 250 worker bees and reproductive queens were established and fed on sucrose only (control) or sucrose solution mixed with dsRNA carrying a foreign GFP sequence (dsRNA-GFP). Control and treated hives were kept in separate net houses. Worker jelly was collected from brood cells containing 5^th^ instar worker larvae and royal jelly was harvested from queen brood cells with 3^rd^-4^th^ instar larvae (see Materials and Methods). We could detect the presence of dsRNA-GFP by Northern blot analysis performed on total RNA extracted from worker and royal jellies. Notably, while the full-length dsRNA could be detected, additional degraded or processed GFP-RNA forms appear in both jellies (Figure 1C).

### RNA is horizontally transferred among bee generations

The presence of ingested dsRNA in the circulatory system indicates systemic spread and the possibility of dsRNA transport to the food glands (Figure 1A, B). This was further supported by dsRNA presence in both worker and royal jellies (Figure 1C). Hence, dsRNA could presumably be transmitted from nurse bees to the young larvae through jelly consumption.

In order to test whether RNA can be transferred horizontally among bee generations down the line, we established reproductive mini-hives that were fed on sucrose solution (control hives) or sucrose solution containing dsRNA-GFP (See Material and Methods). During the experiment, adult workers, larvae, pupae and newly emerged bees were collected and analyzed for the presence of dsRNA-GFP. The experimental design is illustrated in figure 2A. RNA slot-blot assay with a GFP-specific probe was conducted and showed, as expected, the presence of dsRNA-GFP in adult workers that consumed directly dsRNA in their diet. It also demonstrated the occurrence of dsRNA-GFP in larvae that consumed jelly secreted by treated bees (Figure 2B). As previously reported, dsRNA is stable and persists in treated adult bees for a few days post feeding [25]. Here we show that dsRNA, which is transmitted from nurse bees to the larvae, persists in subsequent developmental stages including pupae and newly emerging bees. It appears that ingested foreign dsRNA diminished with time, but could be detected at least 14 days after the last dsRNA feed (Figure 2B).

**Figure 2.**
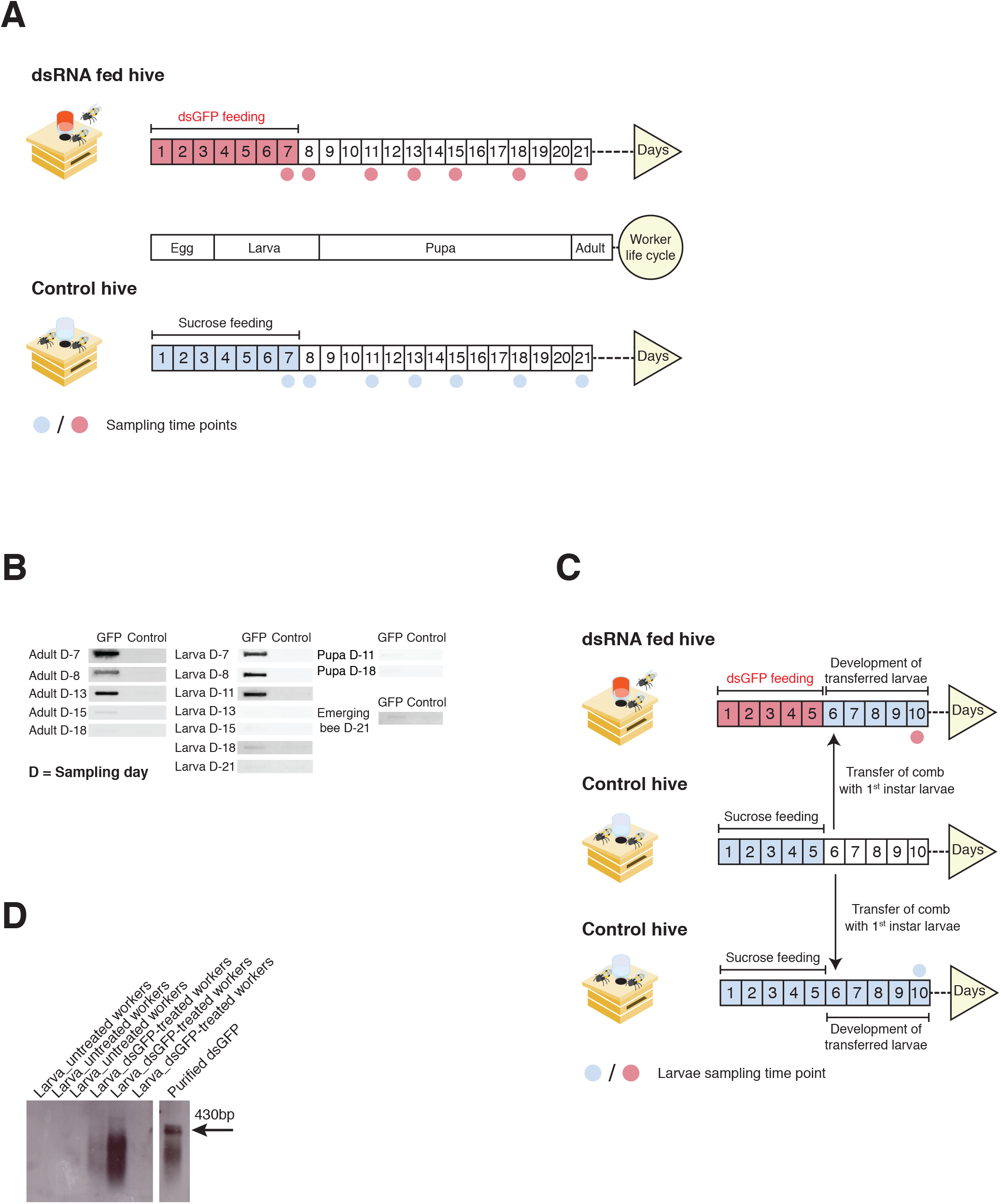
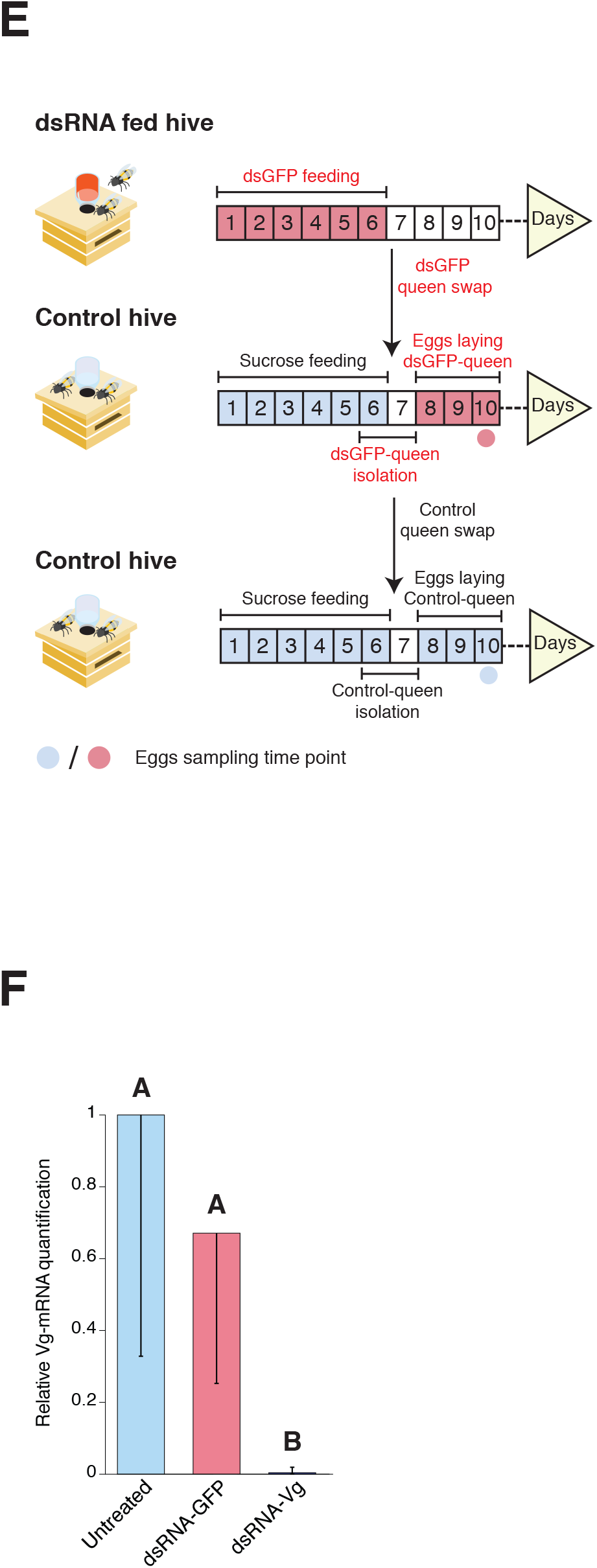
**A)** Experimental design for detecting RNA transmission within the hive. Reproductive minihives contained ca. 250 bees and an active queen. Treated colonies were provided with 300 μg dsRNA-GFP in 50% sucrose solution (w/w) per feeding. Control minihives were fed on 50% sucrose solution (w/w) only. Adult bees and brood were sampled from the available developmental stages. **B)** Occurrence of dsRNA in adult bees and its transfer to the next generation. RNA slot-blot analysis of 1.5 μg total RNA extracted from individual larvae, pupae and adult bees from dsRNA-GFP treated or untreated colonies (Figure 2A). **C)** Experimental design to test horizontal RNA transfer among bees. Reproductive minihives contained ca. 250 bees and an active queen. Treated colonies were provided with 200 μg dsRNA-GFP in 50% sucrose solution (w/w) per feeding. Control minihives were fed on 50% sucrose solution (w/w) only. **D)** Transfer of dsRNA from worker bees to nourished larvae. 1^st^ instar larvae were transferred from untreated hive and nourished by workers from dsRNA treated or untreated minihives for four days. Northern blot of 5 μg total RNA extracted from individual 5^th^ instar larvae. **E)** Experimental design to test vertical RNA transfer among bees. Reproductive minihives contained ca. 250 bees and an active queen. Treated colonies were provided with 200 μg dsRNA-GFP in 50% sucrose solution (w/w) per feeding. Control minihives were fed on 50% sucrose solution (w/w) only. Queens from dsRNA treated or untreated minihives were removed to untreated minihives. The queens were then isolated for two days to acclimatize. Next, the queens were released and their newly laid eggs were collected. **F)** Transmissible RNA is biologically active. Vitellogenin (Vg) knockdown in ten-days old workers that were nourished as larvae by dsRNA-Vg treated bees. Vg expression was quantified by RT-qPCR. Values were normalized relative to a reference calmodulin gene. Individual workers were tested in the untreated (N=6), dsRNA-GFP (N=6), and dsRNA-Vg (N=7) treatments. Data are shown as the mean ±SE. Different letters above the plots indicate statistically significant difference according to the Tukey-Kramer (HSD) test (P<0.05).

While the presence of dsRNA-GFP in the jellies supports horizontal transfer to the progeny, it might also be explained by vertical dsRNA transmission from queen to-egg. To distinguish between the two possible transmission routes, we designed an experiment in which only horizontal nurse bee to-larvae transmission could occur (experimental design is illustrated in Figure 2C). A similar mini-hives set up was established and sucrose solution containing dsRNA-GFP or sucrose solution only (control hives) were provided for five days. On day six, combs containing 1^st^ instar larvae from control hives were transferred either to dsRNA-treated or to another control hive. The transferred larvae were then allowed to develop four additional days in the new untreated- or dsRNA-treated host colonies. On day ten, the 5^th^ instar larvae were collected from the transferred combs, washed rigorously and analysed for the presence of dsRNA-GFP. Northern blot analysis performed on total RNA detected a GFP-RNA signal in larvae that were nourished by dsRNA-treated bees, demonstrating an environmentally mediated horizontal RNA transfer route among honey bee generations (Figure 2D).

RNA transfer from nurse bee to larvae does not rule out paternal queen deposition of RNA to eggs and its persistence throughout the progeny development. Therefore, we attempted exploring whether vertical dsRNA transmission also occurs in honey bees (experimental design is illustrated in Figure 2E). To this end, the reproductive mini-hive system was applied and dsRNA-GFP containing sucrose solution or sucrose only solution (control hive) were provided for six days. On day six, simultaneous queen swaps among treatments were performed as follows: i) the queen from dsRNA-GFP hive replaced the queen from a control hive; and ii) the substituted control queen replaced a queen from a different control hive. On day eight, after two days of acclimatization, the newly introduced queens were released and allowed to lay eggs in their new colony. On day ten, eggs were collected, pooled and analyzed for the presence of dsRNA-GFP. We could not detect GFP-RNA signal in eggs laid by a queen that was previously nourished with royal jelly provided by dsRNA-treated bees (Figure 1C). We acknowledge that such negative detection could be explained by the sensitivity limitation of the Northern blot assay. Thus, we conclude that while vertical transmission might occur to some extent, horizontal nurse to-larvae transfer is the main route to distribute RNA among honey bees.

### Transmissible RNA is biologically active in recipient bees

We next asked whether horizontally transferred RNA (i.e. transmissible RNA) is biologically active within recipient individuals. It has been previously demonstrated that supplementing dsRNA into the natural larval diet induces potent RNAi against endogenous RNA [29,30]. Moreover, similar application with dsRNA, corresponding to Deformed Wing Virus (DWV) and Sacbrood Virus (SBV), effectively reduced viral RNA titter as well as disease symptoms in infected worker larvae and adults [31,32]. Notably, these studies showed that ingestion of jelly-containing dsRNA results in sustainable gene silencing that lasts until adulthood [29–31].

Unlike previous reports, we examined whether dsRNA, which is originated from nurse bees and secreted into the jelly, could elicit RNAi response in the progeny. To answer this question, we established a minimal hive system in plastic boxes containing ca. 150 workers and a comb with eggs and young larvae. The adult bees were fed for eight days on sucrose solution (untreated), sucrose solution mixed with dsRNA-GFP (non-specific dsRNA control) or dsRNA that matched the vitellogenin mRNA sequence (dsRNA-Vg). When the brood cells were sealed, the adult bees were removed and the combs were kept until new workers emerged. We then collected ten days old workers and analyzed the expression levels of vitellogenin by RT-qPCR. Consistent with persistence of jelly-transmitted dsRNA (Figure 2B) and in agreement with the aforementioned reports [29–31], we observed vitellogenin knockdown in adult workers that were nourished as larvae by dsRNA-fed nurse bees (Figure 2F). Therefore, we concluded that transmissible RNA, at least in a dsRNA form, is biologically active in recipient bees.

### Naturally occurring RNA in worker and royal jellies

Royal jelly is a processed food substance secreted from the hypopharyngeal and mandibular glands of young nurse bees. It is mainly composed of proteins, sugars, lipids, vitamins and free amino acids [33]. While queens are exclusively fed on royal jelly, worker larvae consume it only during the first three days post hatching, and then their diet is switched towards worker jelly (mixture of jelly, honey and pollen) [20].

The previous experiments described a mechanism that enables, through jelly consumption, transmission of biologically active RNA among individuals within and between generations in the hive. These findings suggest a naturally occurring transmissible RNA pathway in honey bees. In line with this hypothesis, it has been reported that both worker and royal jellies contain small honey bee RNA populations, demonstrating endogenous bee RNA secretion into the larval diet [17,30]. To further explore the RNA repertoire in the jellies, including the natural occurrence of exogenous and pathogen-related RNA, we adapted a small RNA-seq protocol to sequence full-length RNA up to 200nt (see materials and methods). Samples of royal jelly were collected from 3^rd^ instar queen larvae brood cells, and worker jelly was collected from 5^th^ instar worker larvae brood cells. The jellies were harvested from untreated healthy looking hives.

Size distribution analysis of sequenced RNA indicated that worker and royal jellies have different profiles, with RNAs corresponding to 39- and 72 nt mainly differentiate among the two jellies (Figure 3A, Supplementary Figure 1A). We next applied a metagenomics analysis to identify the origin of jelly RNA and again, found different profiles in the worker and royal jellies (Figure 3B, Supplementary Figure 1B). Surprisingly, bee RNA represents only a minor fraction in both jellies, representing 0.58% and 3.55% of worker and royal jelly, respectively. Instead, large proportions of plant, fungi and bacteria were identified alongside sequences originating from unknown sources. Remarkably, RNA corresponding to various exogenous bee-affecting viruses could also be detected in both jellies.

**Figure 3.**
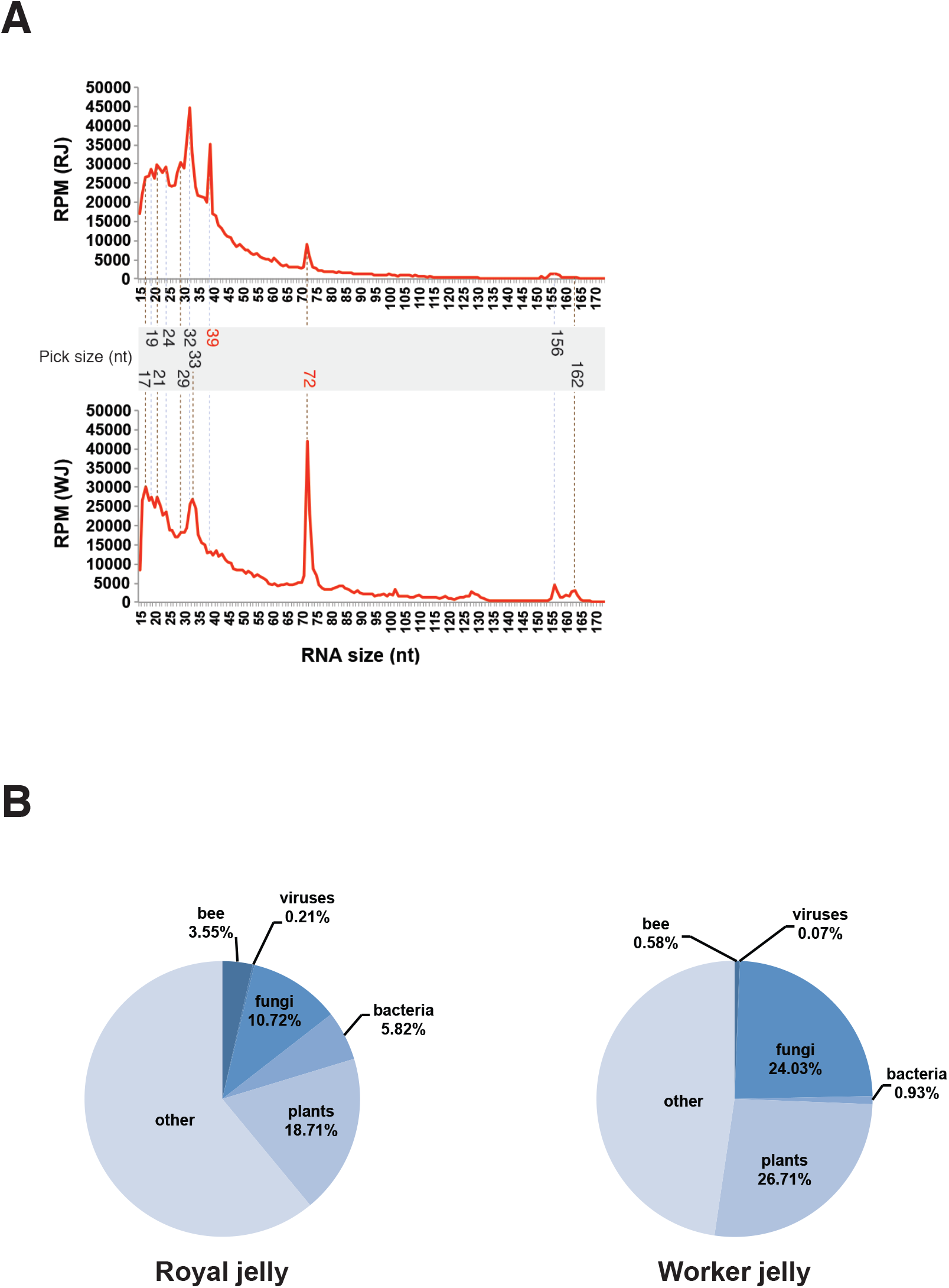
**A)** Size distribution of naturally occurring royal and worker jelly RNA. RNA size was determined through sequencing full-length jelly RNA by a small RNA-seq protocol that was adapted to sequence broad full RNA length spectrum (i.e. 15-200 nt). Data represent a merged analysis of three biological repeats per jelly and are presented as the normalized number of Reads Per Million (RPM). Common peak sizes are marked in black font, and differential sizes in red font. **B)** RNA-based metagenomic analysis of royal and worker jellies. Data represent a merge analysis of three biological repeats per jelly. Individual biological sample analyses are presented in Supplementary Figures 1A and 1B.

We next further characterized jelly RNA that corresponds to the honey bee genome (Figure 4A, Supplementary Figure 2A-C). Like the previous jelly RNA analyses, different honey bee RNA profiles were detected in worker and royal jellies. In both jellies the large proportions represent bee RNA homologous to protein coding genes followed by tRNAs. However, worker jelly is relatively enriched in ribosomal, transposable elements (TE) and non-coding RNA. Interestingly, differential TE RNA occurrence could be detected among the jellies, which is mainly associated with LTR-retrotransposons and TIR transposons (Figure 4B).

**Figure 4.**
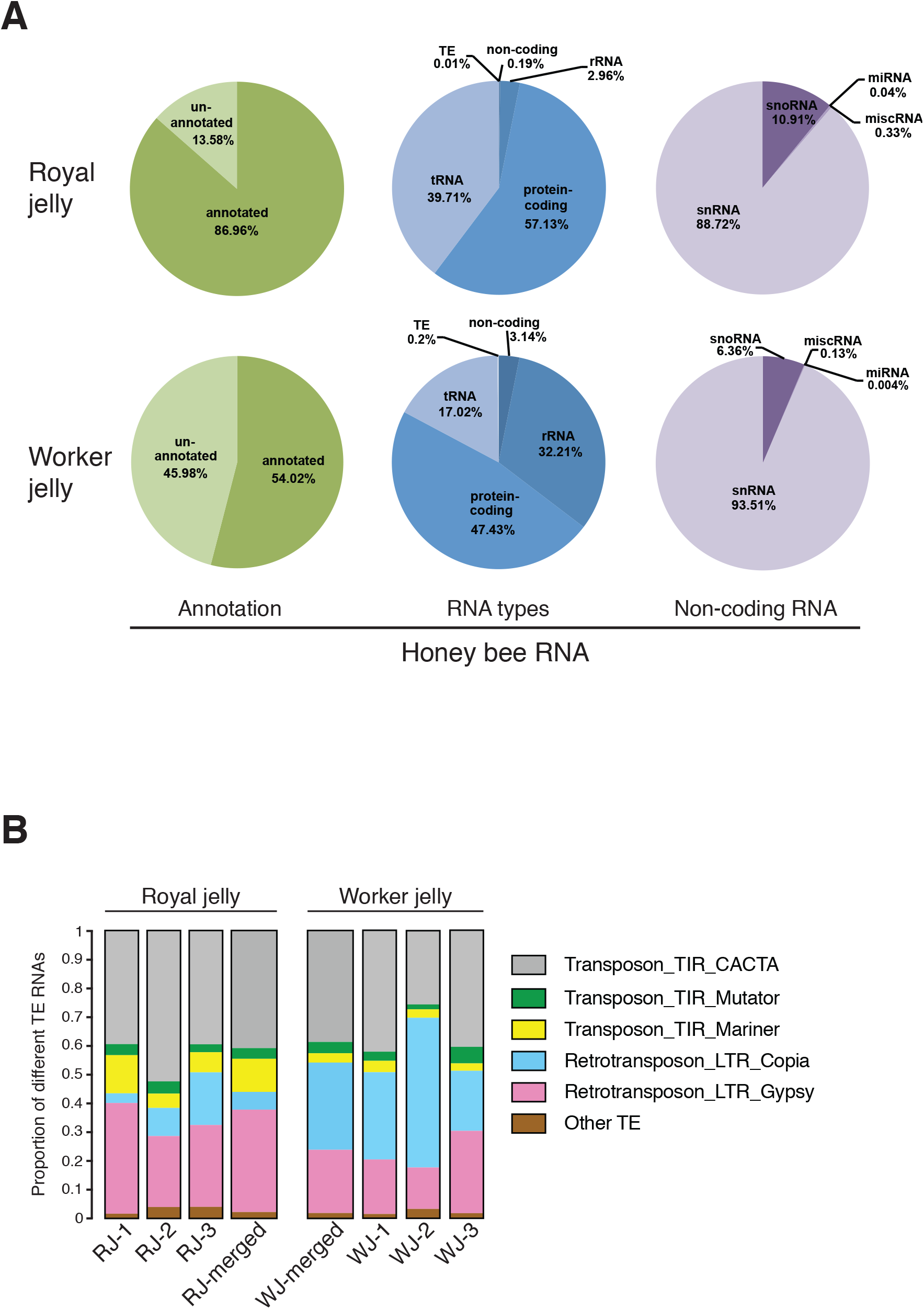
**A)** Classification of honey bee (*Apis mellifera*) RNA in royal and worker jellies. RNA types classification derived from annotated sequences in worker and royal jellies (54.02 % and 86.96 %, respectively). Non-coding RNA classification derived from the RNA types analysis of worker and royal jellies (3.14 % and 0.19 %, respectively). **B)** Occurrence of honey bee transposable elements RNA in royal and worker jellies. TE classification derived from RNA types analysis of worker and royal jellies (0.2 % and 0.01 %, respectively) Data represent a merge analysis of three biological repeats per jelly. Individual biological sample analyses are presented in Supplementary Figure 2A-C

We hypothesized that bees treated with an IAPV-specific dsRNA in previous field trials [26], may have transmitted the antiviral RNA to other bees, resulting in transmissible protection against the viral infection. The presence of naturally occurring viral RNA in both jellies supports the existence of such an RNA-based social immunity in bees. Therefore, we also characterized the viral RNA in worker and royal jellies.

Overall, RNA corresponding to four and ten bee-affecting viruses could be detected in worker and royal jelly respectively (Figure 5A). The most abundant viruses in both jellies were DWV and Varroa Destructor Virus 1 (VDV-1). Interestingly, both sense and antisense viral RNA strands are detected for most viruses. The presence of replicative forms (anti-sense viral genome) suggests an intracellular origin of the viral RNA and its secretion into the jelly rather than RNA derived exclusively from environmental capsids. Additionally, we analyzed the size distribution of viral sequences and identified diverse sense and anti-sense viral RNA fragments in both jellies (Figure 5B). Interestingly, while both worker and royal jellies contain large populations of small RNAs (Figure 3A), almost no small viral RNAs (20-25nt) were identified (Figure 5B). To assess fragment diversity as well as potential occurrence of base-paired viral RNA, reads were mapped against corresponding viral genomes (Figure 5C, Supplementary Figure 3). Multiple mappings were observed in most viruses, especially in the abundant VDV-1 and DWV. Remarkably, presence of long (>25 nt) overlapping viral sense and antisense RNA fragments is somewhat common, suggesting naturally occurring viral dsRNA in worker and royal jellies (Figure 5C, Supplementary Figure 3).

**Figure 5.**
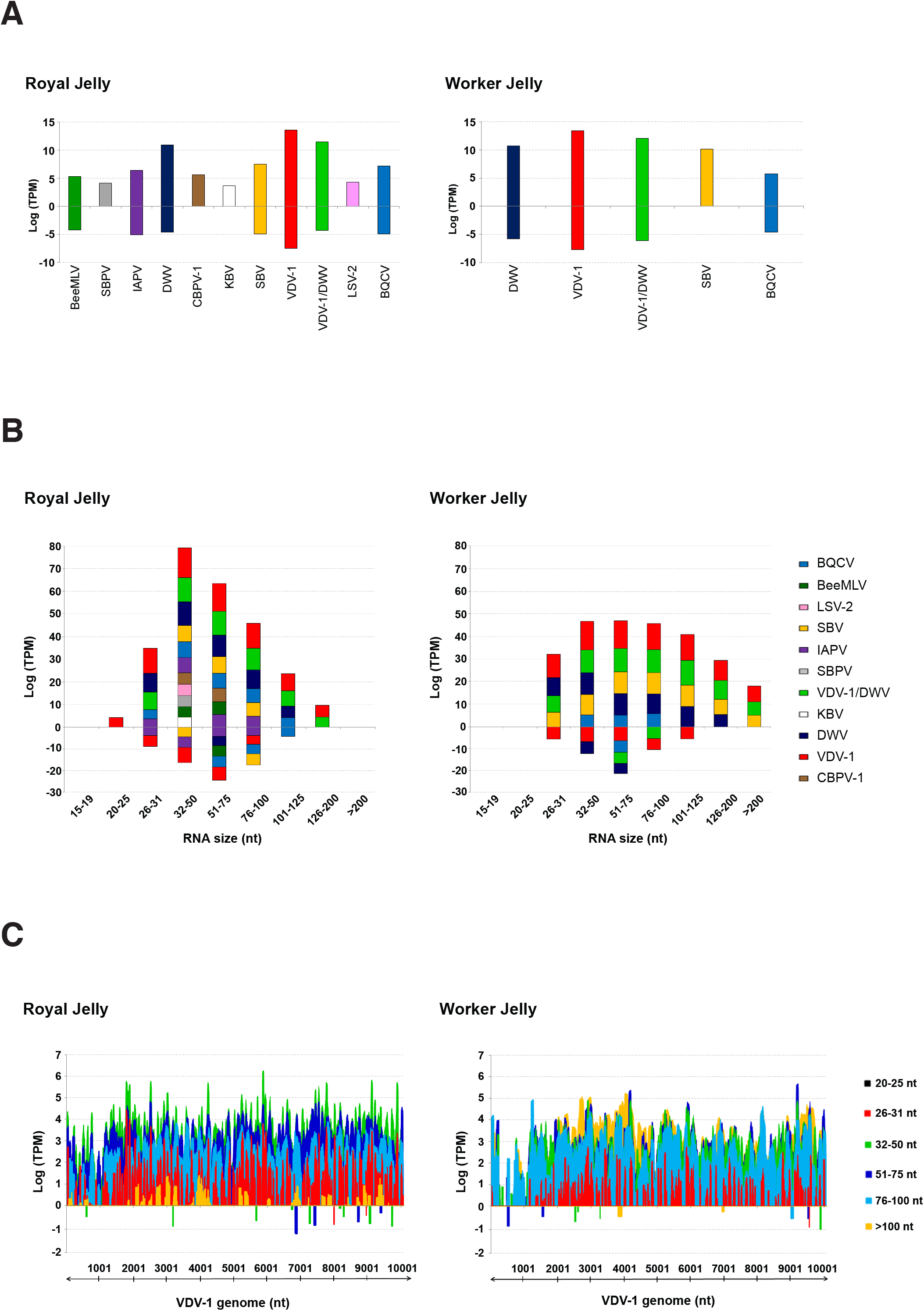
**A)** Natural occurrence of diverse viral RNAs in royal and worker jellies. **B)** The viral RNA in royal and worker jellies is fragmented. **C)** Genome distribution of VDV-1 RNA fragments from royal and worker jellies. Three biological samples were individually sequenced per jelly; sequencing outcomes were merged and analysed. Data are presented as the number of reads normalized to log Transcripts Per Kilobase Million (TPM). Reads that correspond to the sense viral RNA strand (genome) are in positive TPM values, and reads that derived from the antisense viral RNA strand (anti-genome) are in negative TPM values.

## Discussion

Employing dsRNA as a model, this study reports on an environmentally mediated RNA cycle among honey bees. The cycle is engaged by consumption of RNA-containing diet by an individual bee. Then, the ingested RNA is spread from the digestive system through the epithelial gut cells to the hemolymph, where it is associated with a protein complex. A systemic RNA signal reaches the food secretion glands of nurse bees, and is transmitted to the progeny, again, through RNA-containing jelly consumption (Figure 6). This phenomenon is driven by horizontal RNA transfer among individual bees and across generations. Hence, it demonstrates an inherent non-organism autonomous RNA – a transmissible RNA route in honey bees.

**Figure 6.**
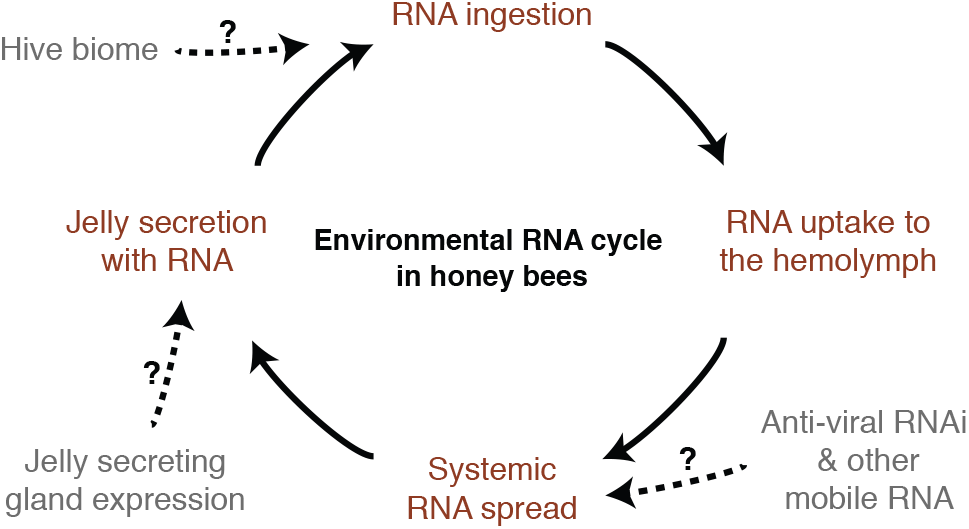
**A)** A working model for transmissible RNA pathway in honey bees. Bees are able to take up RNA from the environment through ingestion. The ingested RNA is taken up from the digestive system through the epithelial gut cells to the circulatory system, the hemolymph. In the hemolymph, ingested extracellular RNA is associated with a protein complex and systemically spread, including to the jelly producing hypopharyngeal and mandibular glands. Then, ingested and other RNAs are secreted in the royal and worker jellies. A new environmental RNA cycle is initiated through ingestion of jelly-containing RNA. A few potential RNA sources could participate in the transmissible RNA pathway including systemic antiviral RNAi and endogenous mobile RNA, jelly-secreting gland transcription as well as hive environmental RNA (e.g. plant, fungi, bacteria).

Larva and adult honey bees can ingest biologically active dsRNA [25,29]. However, the ability of bees to efficiently take up dietary small RNAs (e.g. miRNAs, siRNAs) is currently debatable [17,30,34,35]. While exchange of dietary RNA among adult bees can reasonably occur via trophallaxis [36], in our experiments, secretion of jelly that contained dsRNA-GFP required systemic distribution of the environmentally consumed dsRNA within the nurse bee. A few RNA uptake mechanisms have been reported including internalization by extracellular vesicles, dsRNA channels and receptor-mediated endocytosis [37–40]. Although these mechanisms have demonstrated cellular RNA import, little is known about the systemic spread of RNA through the insect’s circulatory system. Here we show that ingested-dsRNA is exported into the bee’s hemolymph where at least part of it is not naked, but associated with proteins, forming extracellular ribonucleoprotein complexes. In agreement with previous studies, which determined import preference of long dsRNA [39–41] in c. elegans and drosophila, we found that the hemolymph dsRNA-protein complex is comprised of long and mostly non-processed dsRNA (Figure 1B). These findings led us to propose potential parallel complementary functions of the extracellular dsRNA-protein complex involving stabilization, translocation and introduction of the circulatory RNAi signal to recipient cells and tissues in a specific and/or non-specific manner. Yet, investigating such potential roles would require the identification and characterization of the RNA binding hemolymph proteins in future studies.

Suspected prolonged viral disease resistance in field hives fed on dsRNA homologous to a bee virus (IAPV) [26], suggested long-term effect of RNAi in treated colonies 3-4 months after the last dsRNA treatment. RNAi maintenance via dsRNA amplification, driven by the viral RdRp and/or endogenous expression [42–44], can explain silencing persistence in an individual bee. Nevertheless, it is unlikely to explain long-term protection at the colony level since the honey bee’s life span during the summer is ca. six weeks. Worker larvae are fed exclusively royal jelly for three days, and then predominantly jelly, honey and pollen mix. Therefore, the presence of dsRNA-GFP in royal jelly demonstrated jelly-secretion mediated RNA transfer to next generations (Figure 1C). Further supporting such nurse to-larva transfer, dsRNA-GFP could be detected in larvae originated from untreated hives, but nourished by workers fed on dsRNA (Figure 2D). Reciprocally, dsRNA-GFP could not be detected in eggs laid by a dsRNA-treated queen in an untreated hive (Figure 1C). These experiments show that horizontal RNA transfer is the main route to share and spread RNA, at least in a dsRNA form, among the bee population in the hive. However, our results cannot rule out additional vertical transmission of RNA from queen to eggs. Ingestion of dsRNA-supplemented jelly under natural or in vitro conditions has been reported to confer efficient endogenous and exogenous gene silencing in honey bee larvae and newly emerged adults [17,29–31]. Here, we show that dsRNA that is secreted into the jelly and consumed by larvae is also biologically potent and can induce a long lasting RNAi that persists until adulthood (Figure 2B). Therefore, an interpretation of our results leads us to conclude that RNA transfer to larvae could potentially prime anti-viral RNAi and explain the suspected long term protection against viral disease in infected hives fed on IAPV-dsRNA [26].

RNAi has been established in insects, including honey bees, as a key immune response against viruses [25,44,45]. During infection, local RNAi develops into a systemic signal to control viral spread and propagation in distant cells and tissues. Our data indicates that systemic RNAi signal is not limited to the infected bee, but spreads beyond to other individuals in the hive. Diverse fragments of bacterial, fungal and viral RNA naturally occur in both jellies, representing 25% and 16.75% of total worker and royal jelly RNA, respectively (Figure 3B, 5A). The presence of both sense and antisense viral sequences suggests secretion of viral RNA originated from cells. Additionally, potential Dicer substrates seem to be common among natural viral jelly RNA (Figure 5C, Supplementary Figure 3), supporting transfer of naturally occurring RNAi triggers to the larvae. Yet, the relative amount of viral RNA is somewhat low in jelly samples collected from healthy looking hives (Figure 3B). We previously demonstrated cross species bi-directional RNAi transfer between the honey bee and Varroa mite, which has been applied for Varroa control [13,46]. Thus, in addition to individual-defense, transmissible RNA could elicit a colony-level protective outcome. Relative to other insects, the honey bee’s genome encodes a reduced number and variety of immune gene families. It has been therefore suggested that the bees’ behavioral group-defense provides a complementary level of immunity, compensating for the reduction of immune genes [47]. We hypothesize another form of collective defense; social immunity that is engendered through transfer of pathogen-related RNA among members in the hive.

It is generally agreed that RNAi evolved as a defense mechanism against selfish nucleic acids and further diversified to regulate endogenous gene expression. The presence of differential naturally occurring RNA among worker and royal jellies points towards a potential effect of transmissible RNA on genome function in recipient bees. Indeed, supplementing jelly with endogenous or exogenous miRNAs that are naturally enriched in worker jelly affected gene expression as well as developmental and morphological characters of newly emerged workers and queens [17,30]. We speculate that bee to-larva RNA transfer could also play a role in epigenetic dynamics among honey bees. A general involvement of epigenetics in the phenotypic plasticity of female bees has been demonstrated [22–24]. Nonetheless, it is still not understood how caste-specific epigenetic marking is directed. Continuous uptake of regulatory jelly RNA could contribute towards a specific gene-expression profile of a genome with a potential to differentiate into two castes; hence, shifting towards an emergence of worker or queen. Making of a queen is a multilevel process and presumably there are numerous factors interplaying one with the other. Further research should molecularly determine the impact of jelly RNA and its relation to other identified players, such as Royalactin [21].

The presented mechanistic experiments with artificial RNA and the occurrence of natural RNA populations in the jellies indicated not only an RNA share among bees, but also its ability to persist in an external non-sterile environment. It has been reported that royal jelly contains bactericide components [48]. Although inhibition of microbial growth could contribute to the stability of RNAs in the jelly, it cannot solely protect against environmentally distributed nucleases and physical degradation. Moreover, after ingestion, jelly RNAs need to be further stabilized in the digestive system of the larva and adult queen, which harbors a diverse microbial population [49]. This raises the question how does the jelly support environmental persistence and activity of RNA?

## Materials and Methods

### dsRNA synthesis

The GFP sequence and primers as well as dsRNA synthesis procedure were described in Maori et al., 2009. *Apis mellifera* sequence that corresponds to the Vitellogenin mRNA (bases 4648-5084; accession no. NM_001011578.1) served as a template for dsRNA-Vg transcription, and was amplified by the following primers: 5’ **TAATACGACTCACTATAGGGCG**AAAAGCTTATCAGAAGGTGGAAGAAAA 3’; 5’ **TAATACGACTCACTATAGGGCGA**CAATGTTTGTTAACGTTATGGTGGTA 3’ (T7 promoter in bold). Labeling of GFP-dsRNA was performed using a DIG RNA Labeling Kit (Roche) with a DIG-11-UTP concentration of 70 μm per reaction.

### Molecular procedures

Cloning, transcription, RNA preparations, cDNA synthesis, RNA slot blot, Northern-blot, PCR and Proteinase K digestions were carried out according to published protocols (Sambrook & Russell, 2001) or to the manufacturers’ instructions. Northern-blot analyses for the detection of labeled-dsRNA were conducted without using a probe; a standard DIG Northern-blot protocol was modified by omitting the probe-hybridization step. Briefly, 10 μl of pooled whole hemolymph extracts (detailed below) were electrophorased in the Northern-blot analyses of labeled dsRNA, using native 1.2% agarose gel. Slot- and Northern-blot analyses for detection of non-labeled dsRNA were probed with a DIG-labeled PCR probe (Roche Diagnostics Indianapolis, IN, USA) of a sequence corresponding to GFP-sequence used as template for the dsRNA-GFP synthesis.

### Hemolymph extraction from bees

Young worker bees were collected from a single hive and immobilized within plastic straws. Individual bees were fed on 10 μl 50% sucrose solution (w/w) containing DIG-labeled dsRNA-GFP in a final concentration of 50 ng/μl. The control group was fed on 50% sucrose solution (w/w) only. To control dsRNA spillage and cuticle contamination, the immobilized bees were fed directly to their glossa and complete uptake was monitored. Hemolymph was collected from a small incision at the level of the 3rd dorsal tergite, using a microcapillary. The hemolymph of 10 workers was pooled per sample and stored at -80°C for later analysis.

### Reproductive mini-hive system

Caged fertile queen bees, together with approximately 250 worker bees, were placed in mini-hives (26×17×15.5 cm polystyrene) fitted with four mini-combs each. The minihives were sealed and placed in a temperature-controlled room (28°C) for three days in which the combs were constructed and queen-workers recognition had been established. During the first three days, the bees were fed on a mixture of 33% honey and 67% sucrose powder (candy). Next, the mini-hives were transferred into two net-houses separating between dsRNA treated, and untreated mini-hives. The bees were free to fly within the net-houses and to forage for water from buckets. The first 14 days were an adaptation period, during which the colonies were fed on demand with candy, and pollen supplement patties (5 g each) were placed on top of the combs and replaced once a week. An established mini-colony was determined by at-least two constructed combs and egg-laying activity of the queen; only these hives were included in the experiment. During all the experiments, established colonies (two per treatment) were fed on pollen supplement patties (5 g each), and had unlimited water supply.

### Detection of dsRNA-GFP in royal jelly

Treated colonies were fed daily on 10 ml of 50% sucrose solution (w/w) containing dsRNA-GFP in a final concentration of 20 ng/μl (200 μg dsRNA-GFP per day), and untreated mini-hives were fed daily on 10 ml of 50% sucrose solution (w/w) only. Feeding dsRNA-GFP was continued for five days and subsequently the colonies were fed only on 50% sucrose solution until the experiment’s end. At the beginning of day four, the queens were removed in order for the bees to rear new queens. We waited two hours, and placed artificial queen cells containing 1^st^-2^nd^ instar larvae grafted from a different untreated hive. On day 6, we carefully removed intact 3^rd^-4^th^ instar larvae with a fine paintbrush and harvested royal jelly. Royal jelly was harvested from five artificial queen cells, pooled and stored at -80°C.

### Detection of dsRNA-GFP in worker jelly

Treated colonies were fed daily on 10 ml of 50% sucrose solution (w/w) containing dsRNA-GFP in a final concentration of 20 ng/μl (200 μg dsRNA-GFP per day), and untreated mini-hives were fed daily on 10 ml of 50% sucrose solution (w/w) only. Feeding dsRNA-GFP was continued for eight days. On day five and eight, 4^th-^5^th^ instar larvae were carefully removed from worker brood cells and checked for any physical damage. Worker jelly was collected by washing cells with nuclease free water to resuspend the low jelly quantity available. Samples were stored at -80°C.

### Horizontal RNA transfer in the hive

Two treated colonies were fed daily on 15 ml of 50% sucrose solution (w/w) containing dsRNA-GFP in a final concentration of 20 ng/μl (300 μg dsRNA-GFP per day), and two untreated mini-hives were fed daily on 15 ml of 50% sucrose solution (w/w) only. Feeding dsRNA-GFP was continued for seven days and subsequently all the colonies were fed only on 50% sucrose solution until the experiment’s end. The day in which dsRNA was first introduced represents ‘day-1’ (Figure 2A). Samples were collected in different time points, immediately frozen in liquid nitrogen and kept at -80°C for further analysis. Prior to RNA extraction, samples were rigorously washed with nuclease-free water.

### Combs transfer experiment

For five days, treated colonies were fed daily on 10 ml of 50% sucrose solution (w/w) containing dsRNA-GFP in a final concentration of 20 ng/μl (200 μg dsRNA-GFP per day), and untreated mini-hives were daily fed on 10 ml of 50% sucrose solution (w/w) only. On day six, combs containing 1^st^ instar larvae were removed from an untreated hive and transferred either to another untreated or dsRNA-treated colony. On day ten, 5^th^ instar worker larvae were collected, immediately frozen in liquid nitrogen and kept at -80°C for further analysis (Figure 2C). Prior to RNA extraction, samples were rigorously washed with nuclease-free water.

### Queens swap experiment

For six days, one treated colony was fed daily on 10 ml of 50% sucrose solution (w/w) containing dsRNA-GFP in a final concentration of 20 ng/μl (200 μg dsRNA-GFP per day), and three untreated mini-hives were fed daily on 10 ml of 50% sucrose solution (w/w) only. On day six, the queens from both untreated and dsRNA-treated minihives were caged, and on day seven swapped with other queens as follows: the queen taken from the dsRNA-treated colony replaced the queen of an untreated minihive, and a queen from the untreated colony replaced a queen from another untreated minihive (Figure 2E). To allow acclimation, the queens remained caged for a day and then they were released to lay eggs for three days. 100 eggs were collected per treatment on day ten and stored at -80°C for further analysis.

### Vitellogenin silencing by horizontal transfer of dsRNA-Vg

Nine plastic boxes were populated each with ca. 150 workers (collected nearby open brood cells) and a piece of comb containing eggs and young larvae. The plastic boxes were placed in an incubator at 34° C with 45-55% humidity. The boxes were divided into three groups (three boxes per treatment) and were fed daily for eight days with: i) 100μg dsRNA-Vg in 2ml 35% sucrose solution; ii) 100μg dsRNA-GFP in 2ml 35% sucrose solution; and iii) 2ml 35% sucrose solution. Bees were fed with additional 4 ml 35% sucrose solution per day, to avoid hunger. In addition, the bees were routinely fed pollen-sugar supplement to encourage larvae rearing (70% pollen and 30% sugar powder). On day eleven, all cells were sealed and the adult bees were removed. The newly emerged bees were paired for ten days. The pairs were prepared by placing together two bees from different treatments that emerged at the same day and originated from the same hive. Paired bees were fed with 35% sucrose solution and pollen-sugar supplement. On day ten, we collected workers samples from each treatment for Vitellogenin expression analysis.

### Real-time RT-PCR and statistics

qRT-PCR reactions were performed using the EvaGreen^®^ qPCR Plus Mix kit (SOLIS BIODYNE) according to the manufacturer’s instructions. The Vitellogenin mRNA was amplified using primers: 5’-CCAAACTGGAACGGGACCTGC-3’ and 5’- TGTAGCTGTCAGTCGGCGTGC-3’. Calmodulin was used as a control for normalization using primers: 5’-CGAGAGAGAACGGTGGACTC-3’ and 5’- ATACGACACAGCCGACGAG -3’. Statistical analyses were conducted with JMP statistical software version 13 (SAS Institute, Cary, NC, USA). Statistical significance was set at P<0.05. To test for significant differences in relative expression of Vg in workers, one-way ANOVA was conducted on dCt values as previously described [50]. Significant differences between treatments were tested by the Tukey-Kramer (HSD) test.

### Sequencing of full length RNA from worker and royal jelly

Royal jelly was collected from queen brood cells containing 3^rd^ instar larvae, and worker jelly was collected from 5^th^ instar worker larvae brood cells; all brood belonged to untreated healthy looking hives. The jelly samples were immediately frozen in liquid nitrogen and kept at -80°C for further analysis. Total RNA extracted from worker and royal jellies was subjected to Tobacco Acid Pyrophosphatase (TAP) treatment using the Cap-Clip enzyme (CellScript). Modified RNA was purified again by standard Phenol/Chloroform extraction followed by Ethanol precipitation in the presence of Glycogen. The RNA pellet was taken up in 12 ul nuclease-free water. RNA quality and quantity was verified using Agilent RNA 6000 pico chip on a Bioanalyzer 2100 instrument. Libraries construction and sequencing were provided by Cambridge Genomic Services (Cambridge University). Briefly, full-length RNA libraries were prepared using the NEXTflex small RNA-seq kit v3 (Bioo Scientific) according to the manufacturer’s instructions with the following modifications; Adaptor ligation was performed at 16°C overnight (step A). The bead cleanup (step F) was performed following an amended one-sided purification protocol to retain also long fragments (no size selection protocol) as provided by the manufacturer. The final purification of the PCR product (step H1) followed also the amended protocol without size selection as provided by the manufacturer. Sequencing was performed in a NextSeq 500 instrument, in a 150 bp paired-end read run using the high output kit (300 cycle). RNA-seq data have been deposited in the ArrayExpress database at EMBL-EBI (www.ebi.ac.uk/arrayexpress) under accession number E-MTAB-6557.

### Reads trimming and QC

An in-house script utilizing cutadapt (version 1.11) [51], fastx_trimmer (FASTX Toolkit 0.0.13) [52] and FastQC (version v0.10.1) [53] was used to trim raw fastq files. Briefly, Illumina NEXTflex small RNA 5’ and 3’ adapter sequences were trimmed from paired-end fastq sample files, while retaining sequences that were at least 23 bp long. Due to the two colour method of sequencing and other technical sequencing considerations, reads that were shorter than the total number of sequencing cycles had a poly-A tail followed by a poly-G tail that were both trimmed. Then, the four random index bases were trimmed from both ends of the sequences. Next, 5’ and 3’ Illumina NEXTflex small RNA adapter sequences were trimmed anew and only sequences for which the length of both paired reads was at least 15bp long were retained. QC, performed with FastQC, revealed low quality bases at both ends of the reads. These low quality bases were trimmed using fastx_trimmer and an additional QC run indicated that all the samples are properly trimmed. Total number of reads that passed trimming and QC per library is summarized in Supplementary Table 1. All in-house scripts have been deposited in Github and can be downloaded: https://doi.org/10.5281/zenodo.1302437.

### Alignment of royal and jelly samples to the genome of *A. mellifera*

The genome of *A. mellifera* was downloaded from ensembl (release 32). Using in-house scripts (https://doi.org/10.5281/zenodo.1302437), the fastq files (R1 & R2) of each royal and jelly sample were converted into fasta files, which were then aligned to the *A. mellifera* genome using blat [54]. The best hit for each R1 mapped read was matched with the best hit for its R2 mapped read. Therefore determining the final mapping of the read as well as the size of its matching RNA.

### RNA size analysis

For each royal and worker jelly sample, we computed the length of the sequenced RNA’s independent of any genome alignment. That is, we used blat to align an R1 read and the reverse complement of its matching R2 read. We then calculated the length of the sequence the two reads span from the 5’ of R1 to the 3’ of R2. The size of reads that did not overlap is indicated as being greater that the length of the longer between the R1 and R2 reads (e.g. >38 where 38 is the length in bp of the longer read). Using the described method, we managed to map the size of approximately 98.5% of the all the reads in each of the samples

### Comparative RNA size analysis

To compare the size distributions of RNA in royal jelly and worker jelly samples, we computed a histogram of all RNA sizes present in the jellies and calculated the log of Reads Per Million, hereof denoted as RPM, using the following formula:

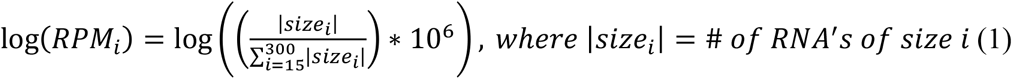

To compute the log(RPM) of royal and worker jelly samples in general, we calculated the average number of RNA’s in each size group for all royal / worker jelly samples, that is 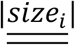, and used it to calculate the general royal / worker log(RPM) value using the following formula:

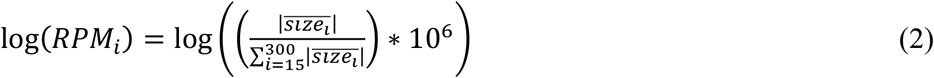

*where* 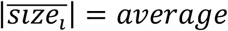 # *of RNA’s of size i in royal or worker jelly samples*

### Comparative metagenomics of sequenced RNA populations

To identify the origin of royal and worker jelly RNAs we used blat [54] to map the RNA sequences against the genomes of *A. mellifera*, 13 viral genomes, 5243 bacterial genomes, 1038 fungi genomes and two plant genomes. The names and the sizes of the genomes are depicted in Supplementary Table 2. Each of the royal and worker jelly RNA samples were mapped to the above mentioned genomes and the percentage of RNA mapped to each specie were recorded as the number of RNA sequences mapped to each specie out of the total number of RNA sequences in the sample. The RNA in each sample were first searched against the bee genome and then against viral genomes, bacterial genomes, fungi genomes and finally against plant genomes. Each blat search was performed with the RNAs that did not match the previous genome they were searched against. That is, all the RNAs in each sample were searched against the bee genome but only RNA reads that did not match the bee genome were searched against viral genomes, only RNAs that did not match any of the viral genomes were searched against bacterial genomes etc’. In order to calculate the number of RNA sequences mapped to each genome for each type of jelly (i.e. royal / worker), the data from the three royal jelly samples were treated as a single sample and the data from the three worker jelly samples were treated as a single sample. We then recorded the percentage of RNAs that were mapped to each species for the merged royal jelly samples and for the merged worker jelly samples. For individual sample analysis, we simply recorded the percentage of RNAs that were mapped to each species for each royal or worker jelly sample.

### Characterization of jelly RNA corresponding to the honeybee genome

We used the ensembl annotation file for the genome of *A. mellifera* (release 32) to characterize the jelly RNA corresponding to the honey bee genome. Using an in-house script, we converted the ensembl annotation file into a bed format file and intersected it with a bed file version of the RNAs corresponding to the honey bee genome using intersectBed (bedtools version 2.17.0) [55]. The number of RNAs in each annotation category was then recorded, excluding matches in which the overlap between the RNA and the annotation element was shorter than 7bp. To further characterize the jellies corresponding to the honey bee genome, we performed a blat search against two transposable elements (TE) databases, the TREP database (release 16) [56] and RepBase (version 22.04) [57]. The number of RNAs that mapped to each of the TE categories was then recorded from which the percentage of each TE category was calculated.

### Detection of viral RNAs

To detect the presence of pathogen-related RNA, we performed a blat search [54] against 13 honey bee viral genomes (Supplementary Table 2). That is, the RNAs of each of the six jelly samples were searched against each of the viral genomes and their genomic position, length (i.e. insert length), orientation (i.e. forward/reverse) and abundance, relative to each viral genome, were obtained (Supplementary Tables 3 and 4).

### Abundance comparison of viral RNAs between royal and worker jellies

To enable a comparison between the abundance of virus-related RNAs in each of the 13 viral genomes, we calculated for each jelly (i.e. royal or worker) its log(TPM), where TPM = Transcripts Per Million, using the following formula:

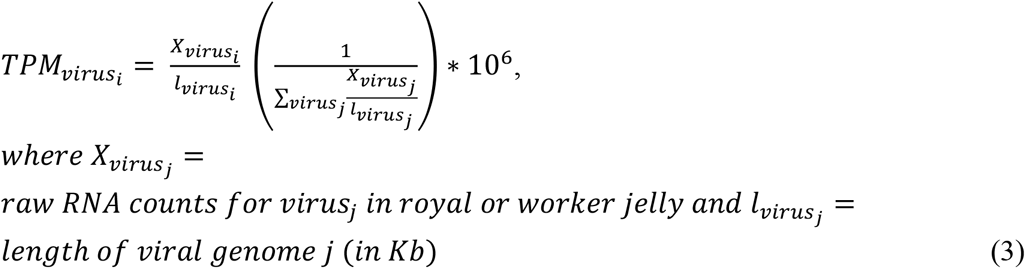

To analyze the size distribution of viral RNAs for each jelly type (i.e. royal / worker) or for each sample separately, we grouped the various insert lengths into nine distinct size groups: 15-19nt, 20-25nt, 26-31nt, 32-50nt, 51-75nt, 76-100nt, 101-125nt, 126-200nt and >200nt and calculated the log(TPM) of each viral genome in each size group using the following formula:

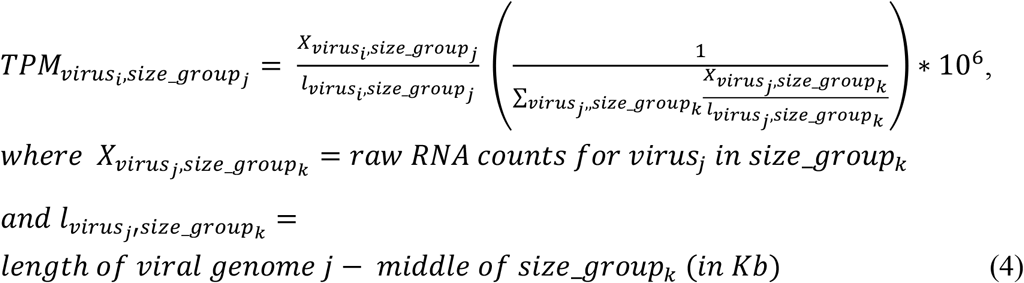

To evaluate the abundance of viral RNAs along the genome of each virus, we calculated the log(TPM) at each position along the viral genome of each virus using the following formula:

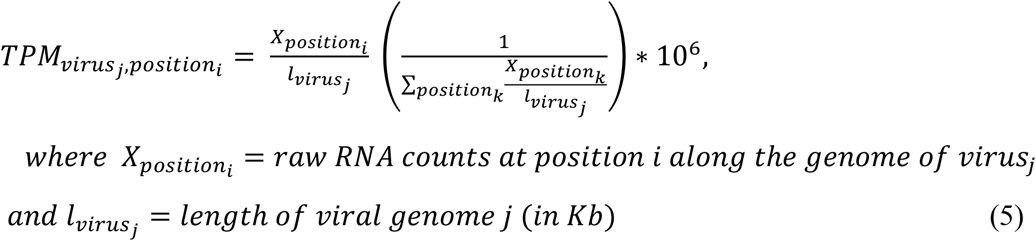

---

**Supplementary Figure 1.**
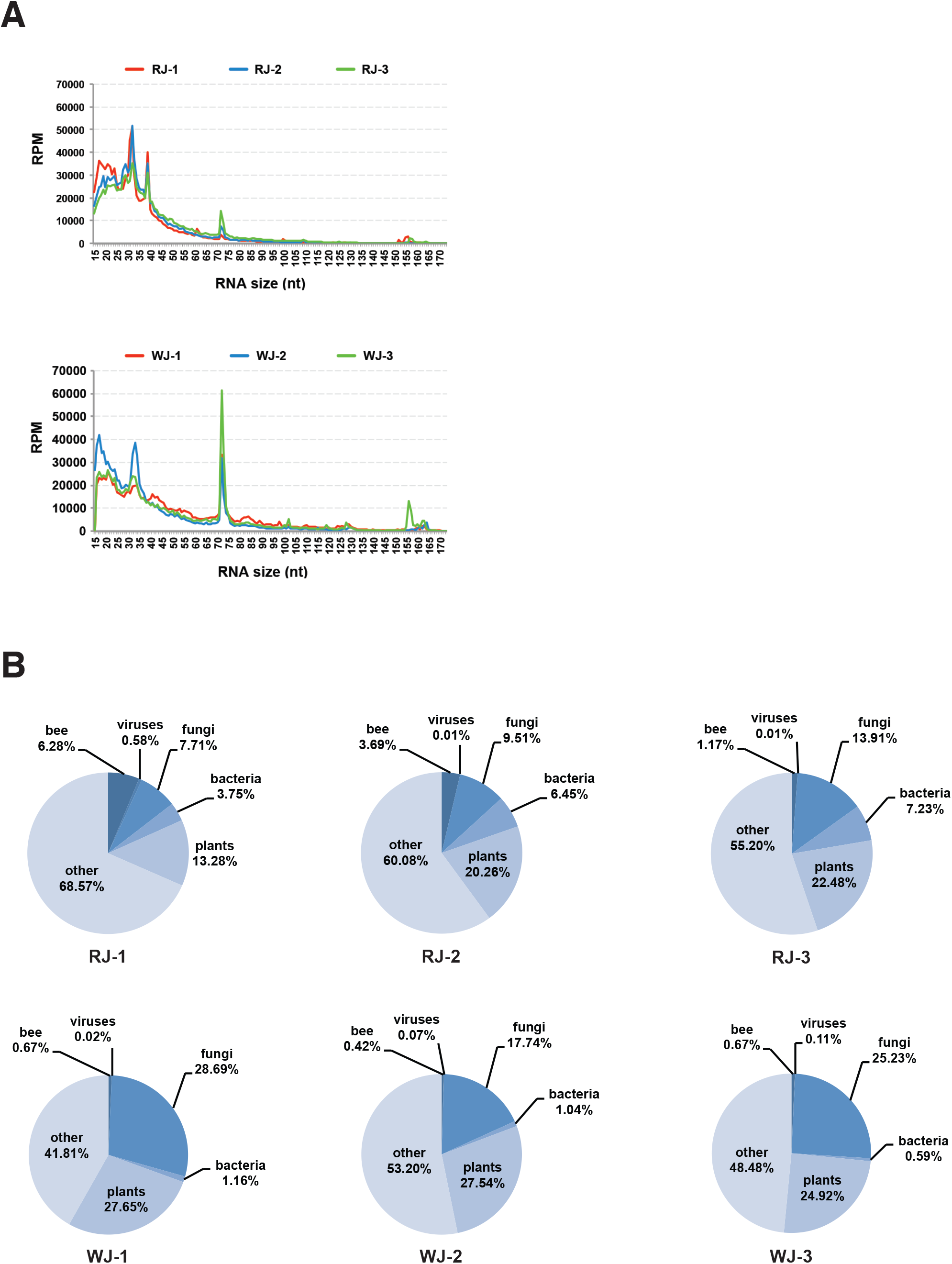
**A)** Size distribution of royal and worker jelly RNA. Data are presented as the normalized number of Reads Per one Million (RPM). **B)** RNA-based metagenomic analysis of royal and worker jellies.

**Supplementary Figure 2.**
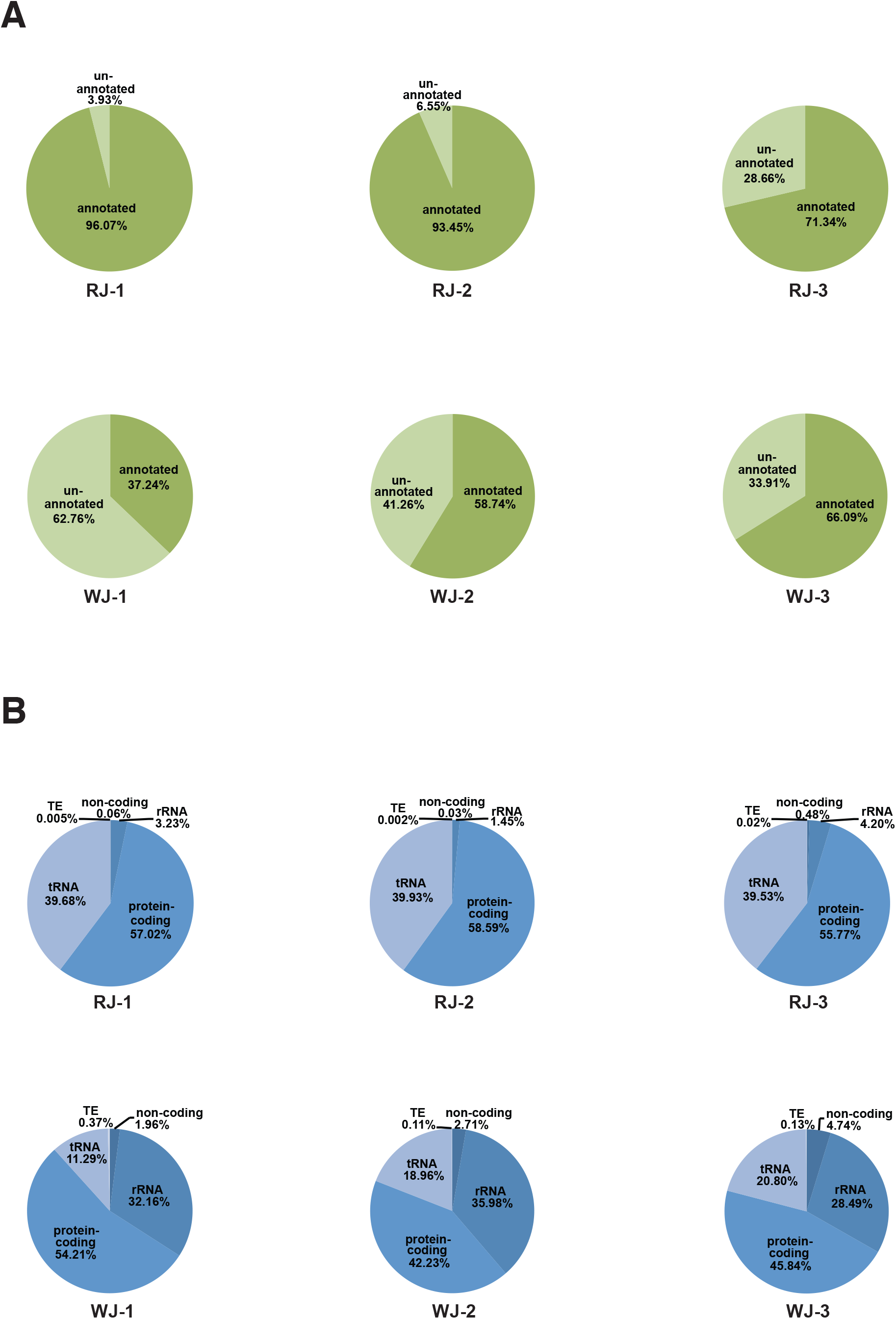
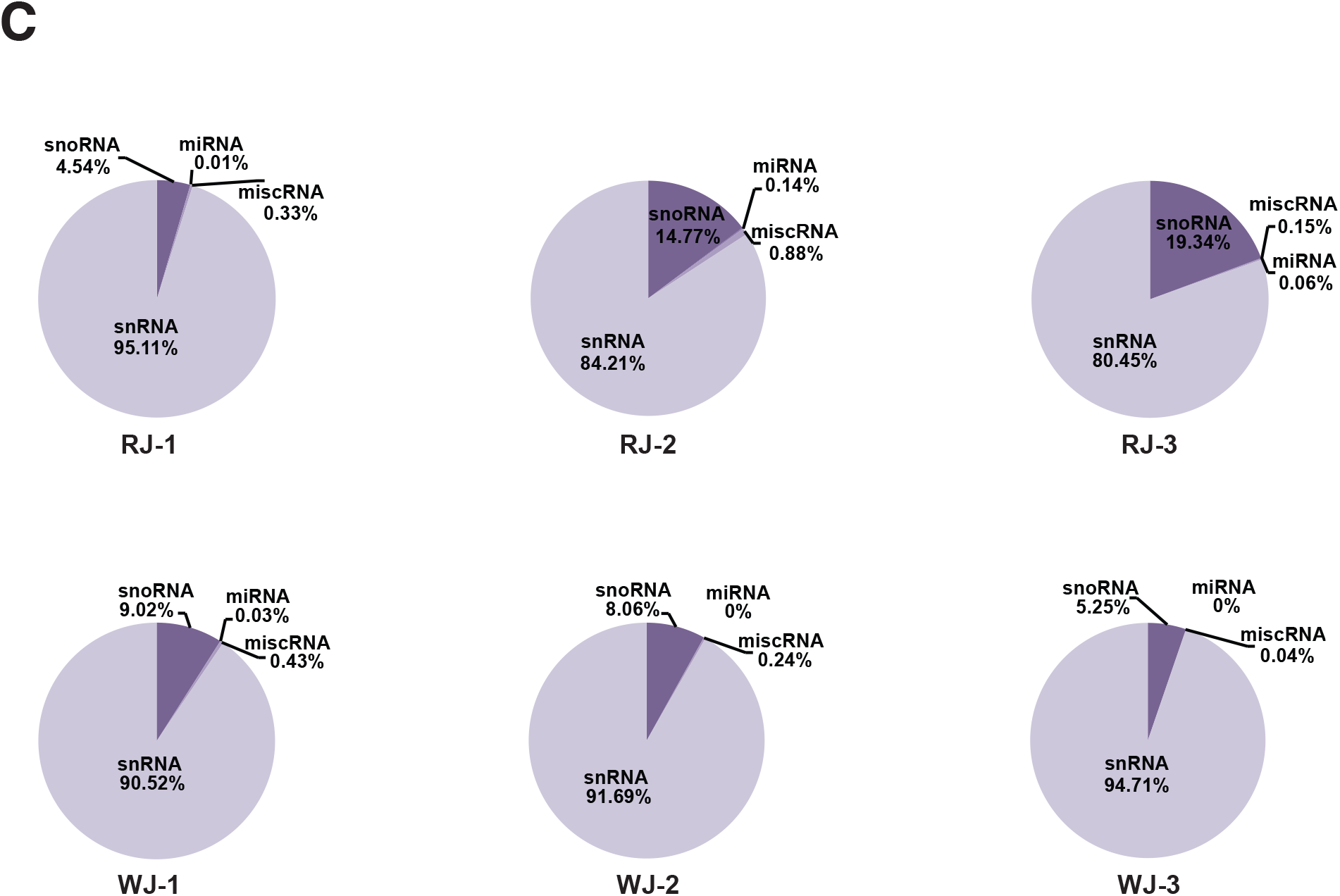
**A)** Proportion of annotated and un-annotated honey bee RNA in three biological replicas of royal and worker jellies. **B)** Proportion of different honey bee RNA species in three biological replicas of royal and worker jellies. **C)** Proportion of non-coding honey bee RNA species in three biological replicas of royal and worker jellies.

**Supplementary Figure 3.**
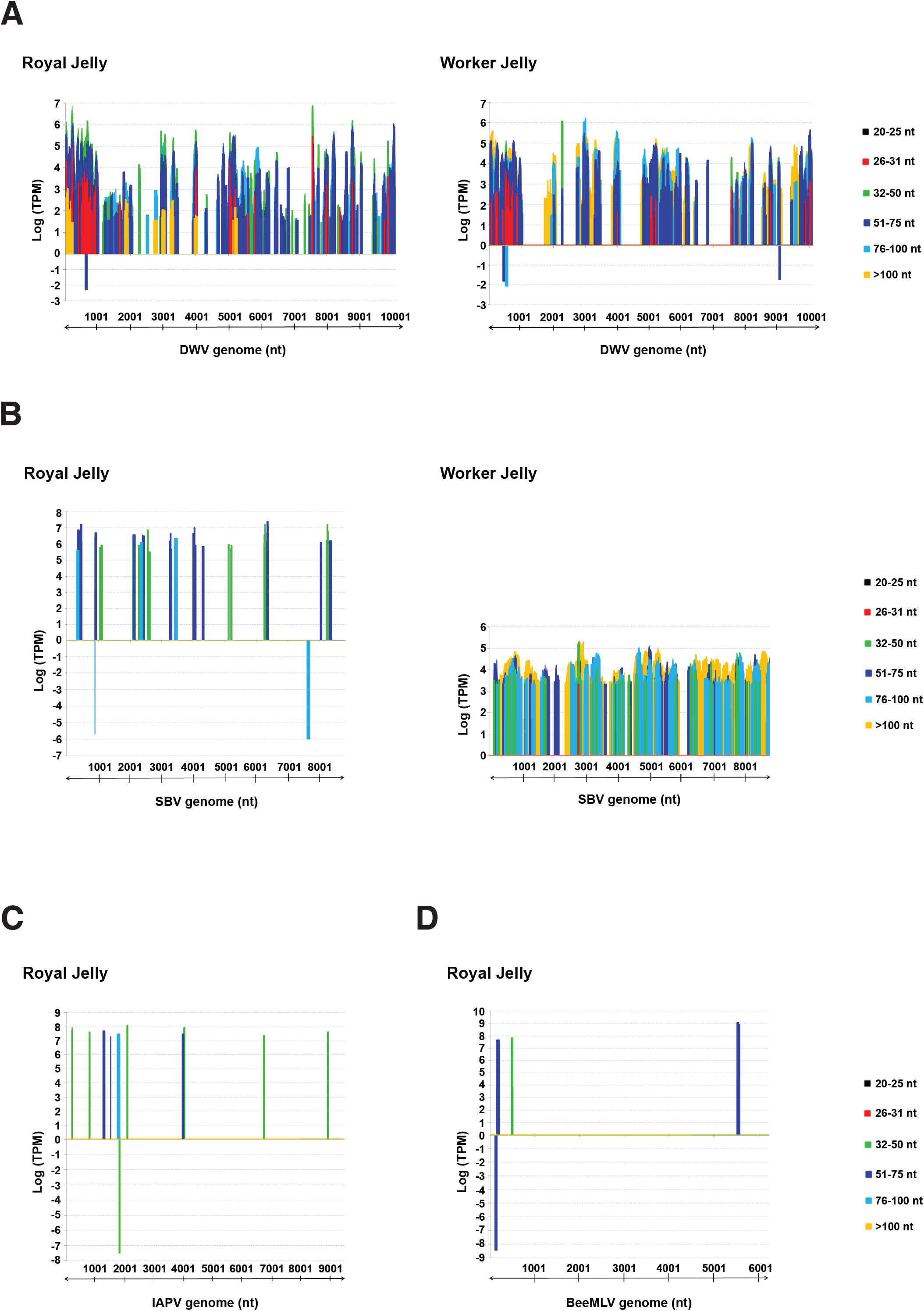
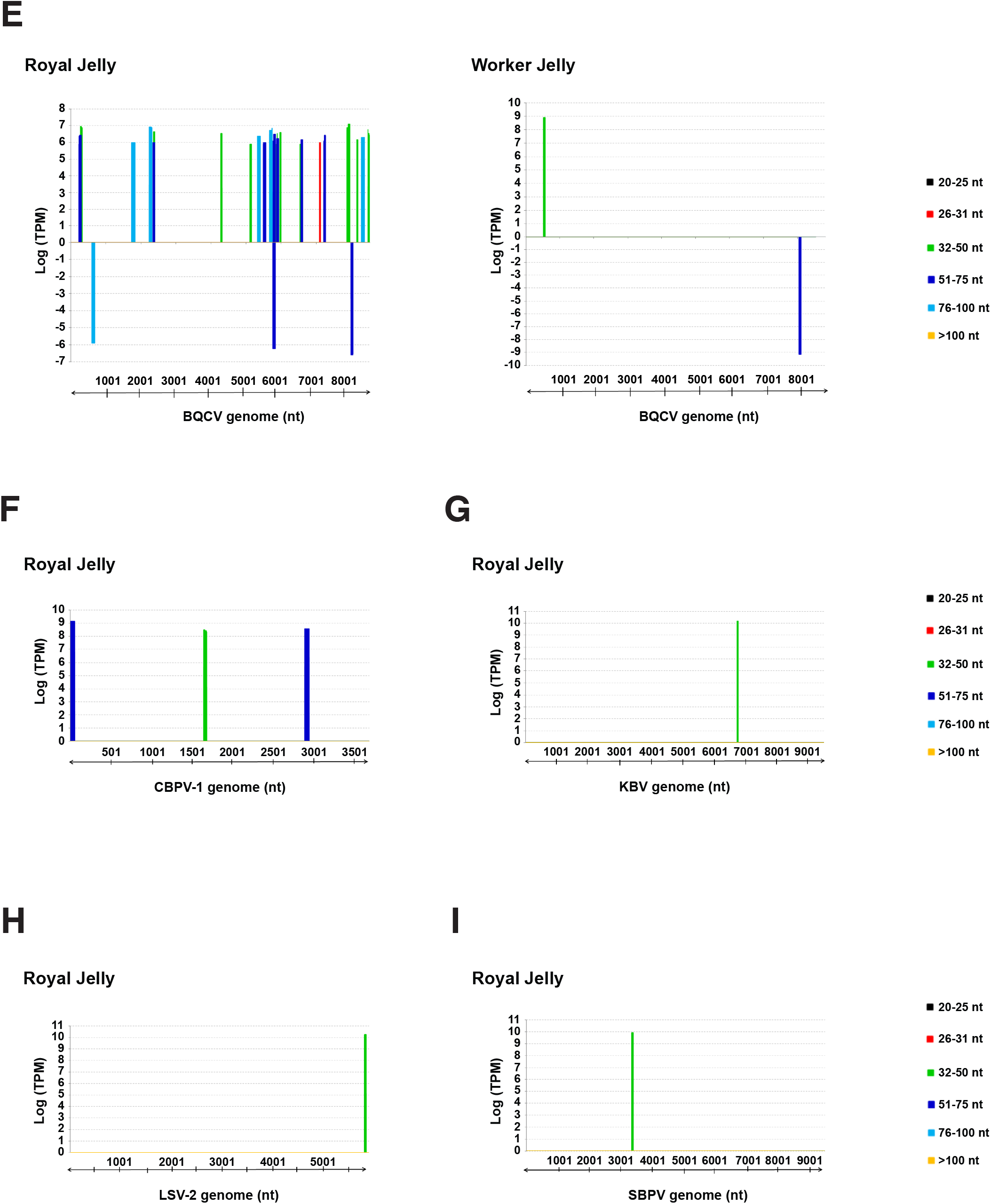
**A)** Genome distribution of DWV RNA fragments from royal and worker jellies. **B)** Genome distribution of SBV RNA fragments from royal and worker jellies. **C)** Genome distribution of IAPV RNA fragments from royal jelly. **D)** Genome distribution of BeeMLV RNA fragments from royal jelly. **E)** Genome distribution of BQCV RNA fragments from royal and worker jellies. **F)** Genome distribution of CBPV1 RNA fragments from royal jelly. **G)** Genome distribution of KBV RNA fragments from royal jelly. **H)** Genome distribution of LSV2 RNA fragments from royal jelly. **I)** Genome distribution of SBPV RNA fragments from royal jelly. Three biological samples were individually sequenced per jelly; sequencing outcomes were merged and analysed. Data are presented as the number of reads normalized to log Transcripts Per Kilobase Million (TPM). Reads that correspond to the sense viral RNA strand (genome) are in positive TPM values, and reads that derived from the antisense viral RNA strand (anti-genome) are in negative TPM values. Not all viruses could be detected in both jellies.

**Supplementary Table 1**.

Worker and royal jellies RNA-seq read counts after trimming and QC

**Supplementary Table 2**.

List of species used in the metagenomics analysis

**Supplementary Table 3**.

Summary of viral sequencing reads obtained from royal jelly sequencing

**Supplementary Table 4**.

Summary of viral sequencing reads obtained from worker jelly sequencing

## Acknowledgments

Professor Ilan Sela, who has died aged 81, was Professor of Virology and Molecular Biology at the Hebrew University of Jerusalem and was regarded as one of the pioneer virologists in Israel. Ilan was a creative mind and a passionate scientist. He found joy in understanding fundamental principles in biology and reveled in their translation into real world solutions. For more than five decades he nourished generations of students with curiosity and admiration towards science. Ilan is greatly missed. The authors thank Dr. Ed Farnell and Dr Emily Clemente from Cambridge Genomic Services for their contribution in the RNA-seq protocol development. We thank Mr. John Rayner (Springwell Apiaries, UK) for his support on bee and jelly samples. We are grateful to Prof. Eric Miska, Prof. Jonathan Heeney and their groups as well as to Professor Sir Peter Lachmann, Mr. David Seilly, Dr. Laurence Tiley and Dr. Shahar Avin (Cambridge University) for inspiring discussions and advice. This work was supported by internal B. Triwaks Bee Research Center funds (HUJI no. 0356043), the Orion Foundation (HUJI no. 0368598), and Israel Science Foundation Grant 1456/10 to SS. Research was also supported by Marie Curie Intra-European Fellowship For Career Development (PIEF-GA-2010-274406), Leo Baeck Scholarship and Herchel Smith Postdoctoral Fellowship (XXACC_AFGLTRB2) to EM.

